## Supplementary Materials (Figures and Tables) for "Imputation of label-free quantitative mass spectrometry-based proteomics data using self-supervised deep learning"

### Supplementary Figures

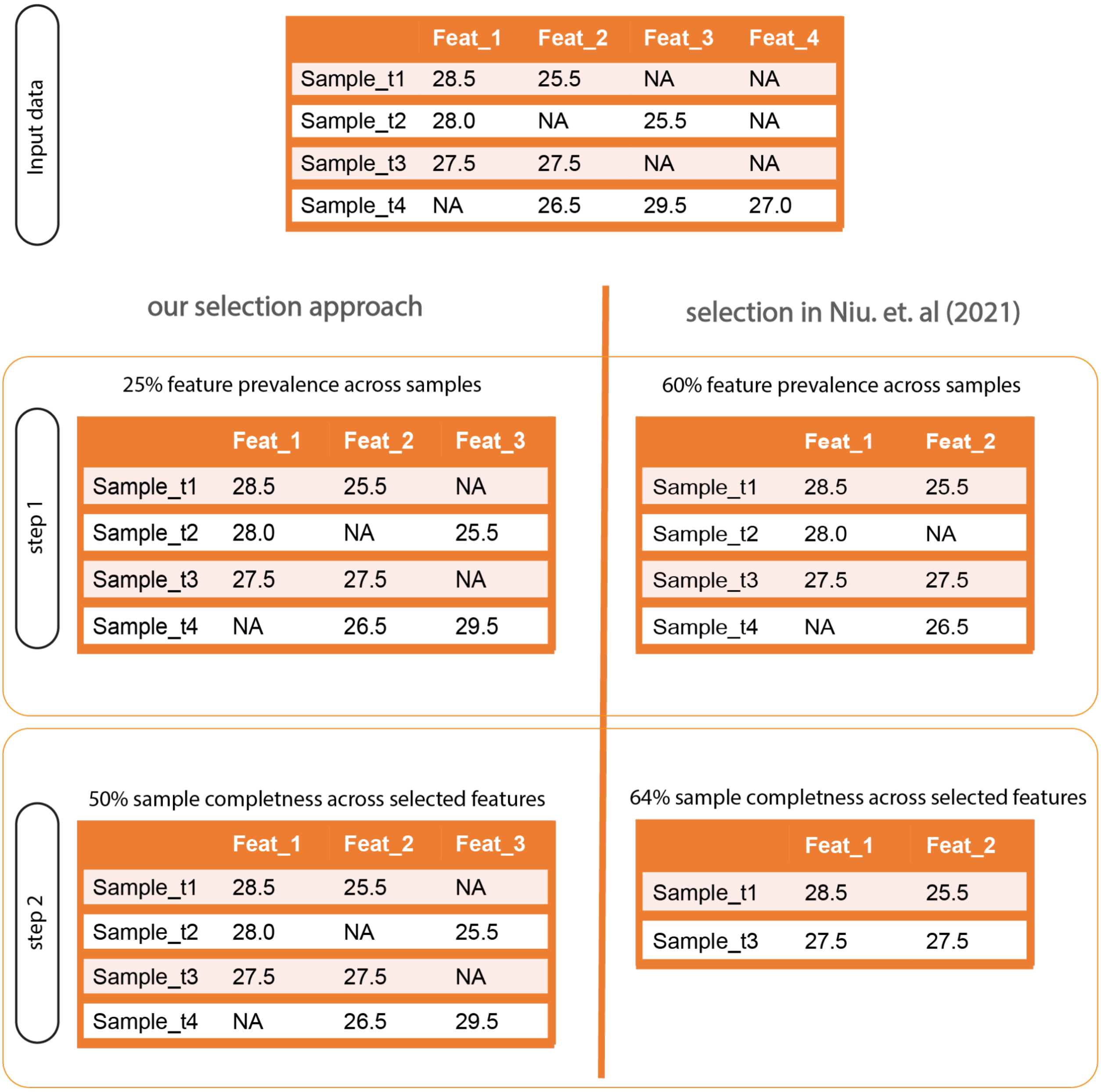


**Fig S1: Comparison between approaches used here and random shifted normal (RSN) imputation for ALD data.** First, features are independently selected by their completeness over a set of samples. Second, samples are selected over the set of selected features from the first step. As we were aiming to impute as many features as possible, we lowered the threshold below 60% in the first step. However, we needed enough observations from a single feature to be able to learn the dependence on other features which worked with a threshold of one quarter in larger data sets (which can be adapted). The second step ensures that samples have some coverage over the prevalent features, i.e. low quality samples get removed, as imputation is done based on other features in a sample.

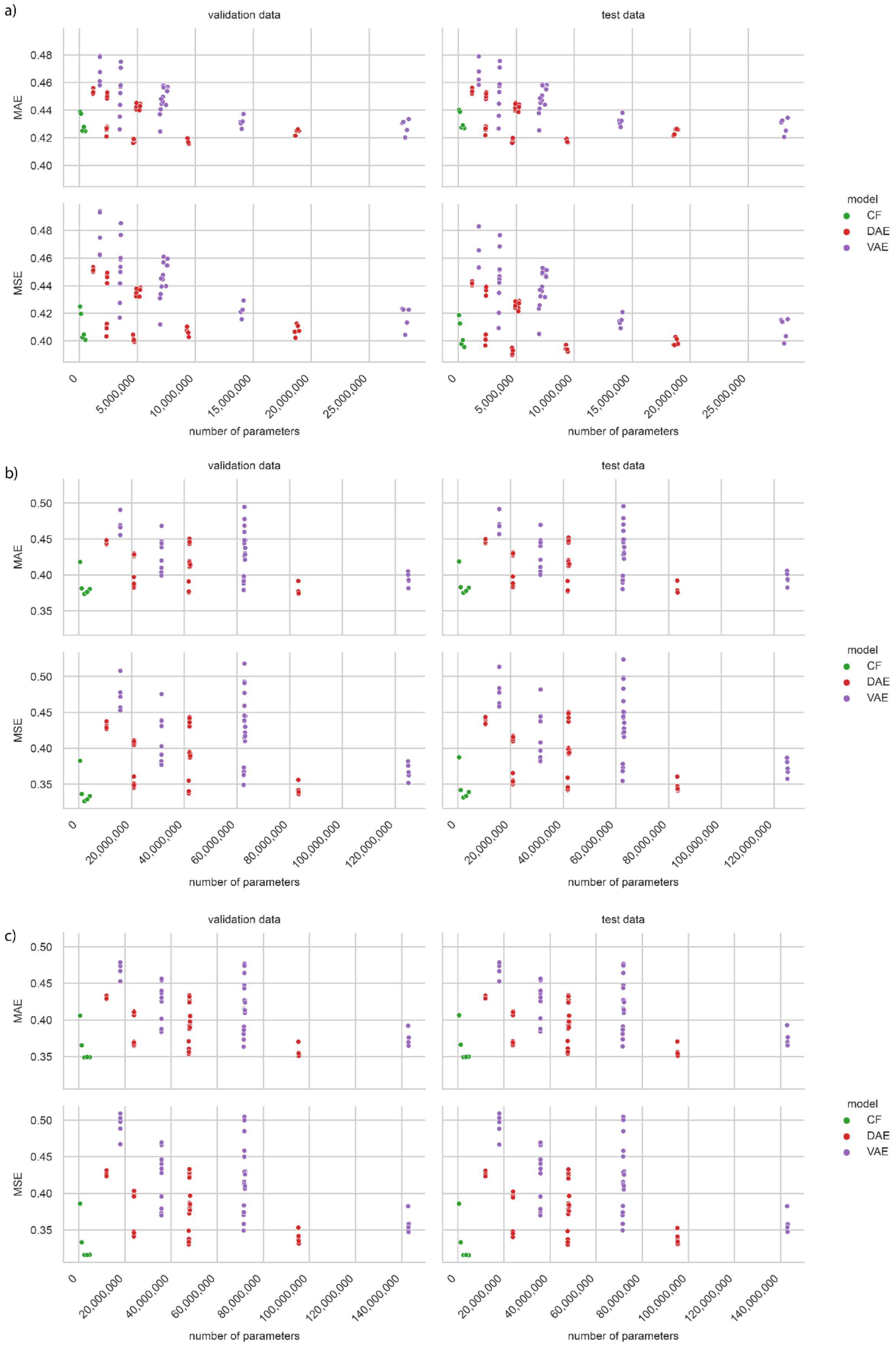


**Fig. S2: Grid search performances by number of model parameters.** The total number of parameters depends on the architectures (which are the same, see Methods) and the number of features. CF performs more stable than DAE and VAE which show more variation in their performance for a) PGs b) aggregated peptides c) precursors. Mean average error (MAE) and mean squared error (MSE) of simulated missing values for both validation and test data.

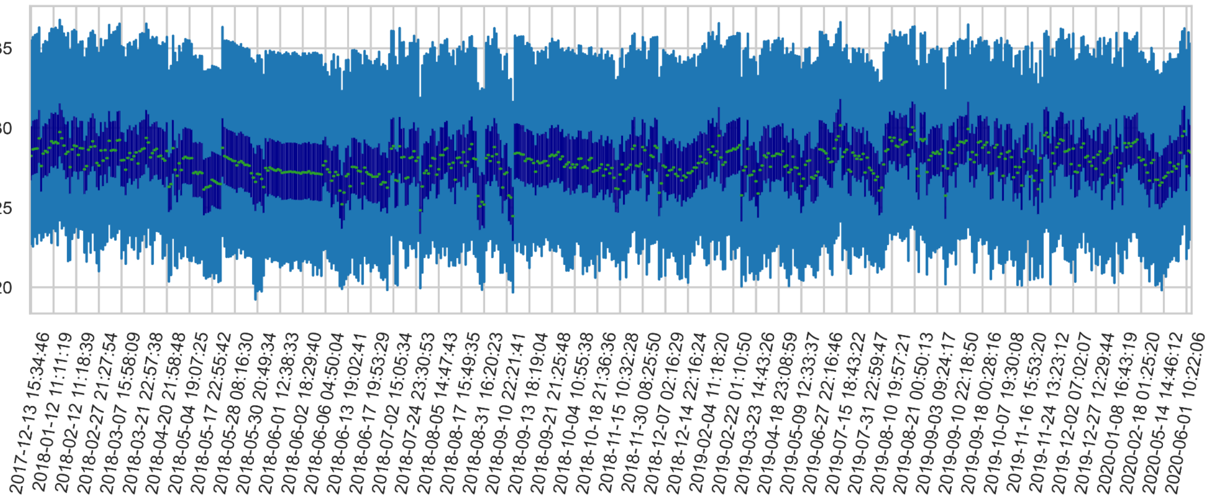


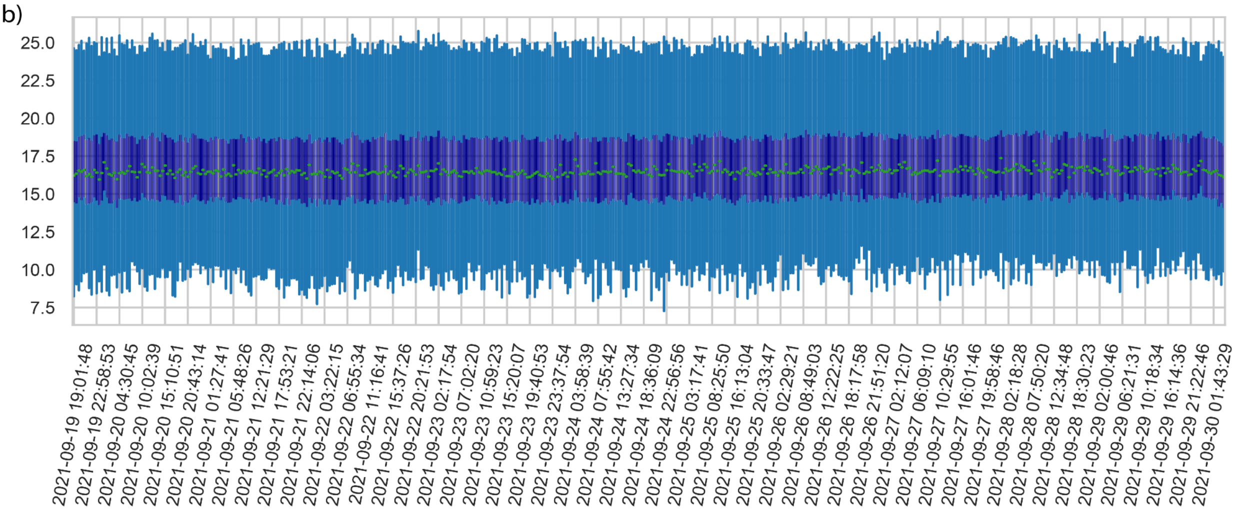


**Fig. S3: Boxplots of each of the HeLa samples for instrument 6070 and ALD plasma sample dataset a)** Boxplot of logarithm two transformed intensities for protein groups (y-axis) of HeLa data for instrument 6070. Notice the long time range of samples and that intensities range roughly from 20 to 36. **b)** Boxplot of logarithm two transformed intensities (y-axis) of plasma samples measured in comparison to a) short time range of several weeks. The values range from 8 to 26. The lower and upper hinges (dark blue) of the boxplots correspond to the first and third quartiles. The upper and lower whiskers (light blue) extend from the hinge to the highest and lowest values, respectively, but no further than 1.5× interquartile range from the hinge. Data beyond the ends of whiskers are outliers and are plotted individually.

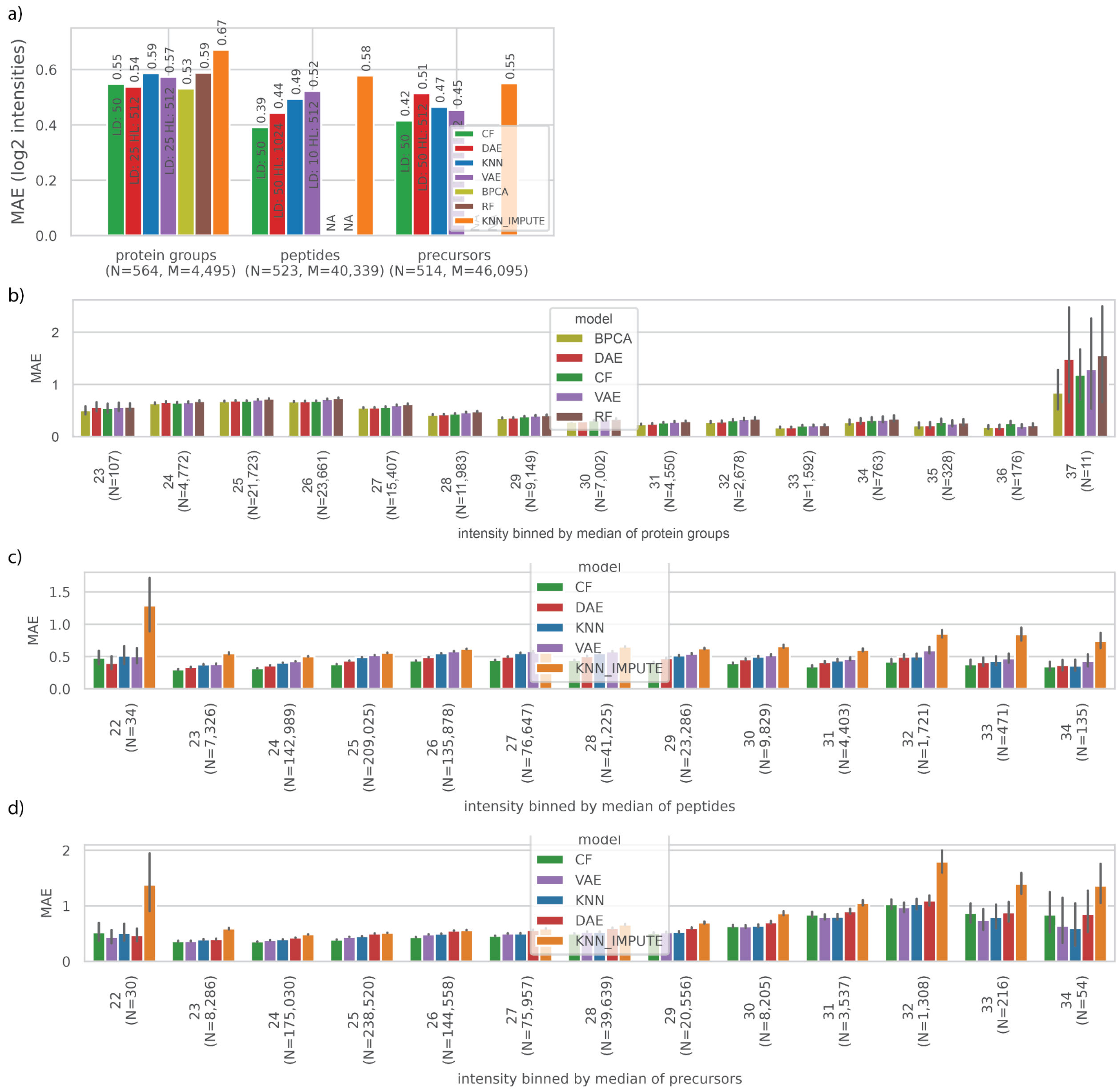


**Fig. S4: Performance for larger dataset for best models according to grid search. a)** Performance of imputation methods at the level of protein groups, aggregated peptides, and precursors. Mean averaged error is shown on the y-axis (green: CF, red: DAE, blue: KNN (scikit-learn), purple: VAE, olive: BPCA, brown: random forest (missForest), orange: KNN_IMPUTE (impute)). The ML methods performed competitively in comparison to the other models. **b)** MAE for protein groups intensities in test split binned by the integer value of their median intensity in the training data split (4,495 protein groups, based on 103,902 intensities, models ordered by overall performance) **c)** as b) for peptides (40,399 peptides, based on 652,968 intensities, models ordered by overall performance) **d)** as b) for precursors (46,095 precursors, based on 715,896 intensities, models ordered by overall performance)

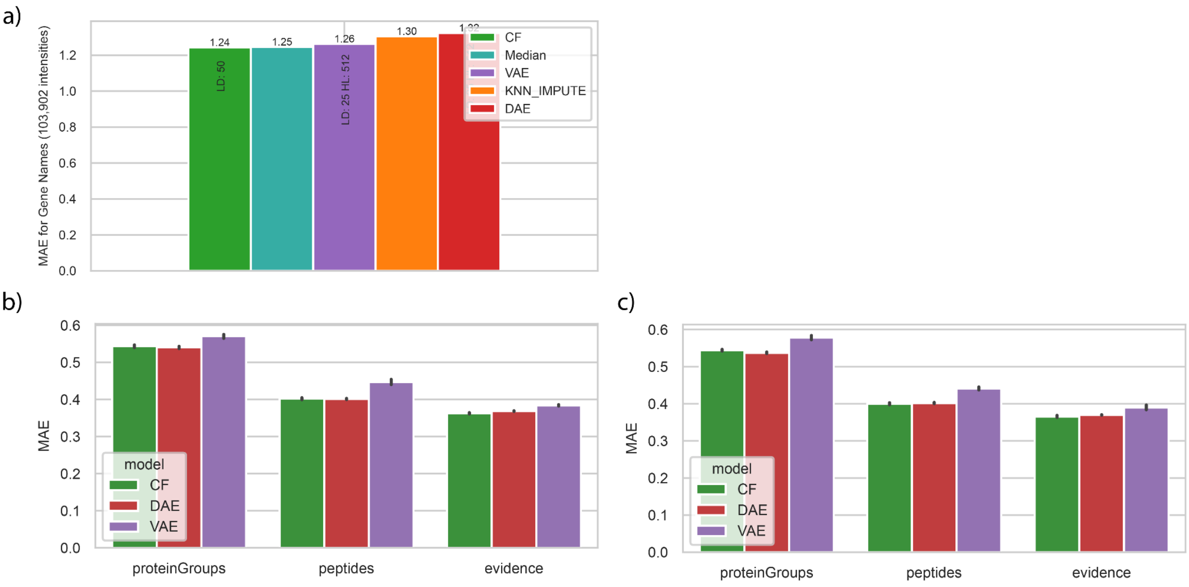


**Fig. S5: Overfitting analysis on large development dataset a)** Fitting large development dataset to randomly permuted protein groups. Median performance is not changed due to permutation and will be close to the best performance on the permuted data. **b)** Repeated training of the best self-supervised deep learning models on different data splits and **c)** repeat training only on the same random split of the data. In both cases the overall performance is stable. All analysis was done with a share of 25 percent MNAR in simulated missing values. See **Supp. Data. 5** for summary statistics.

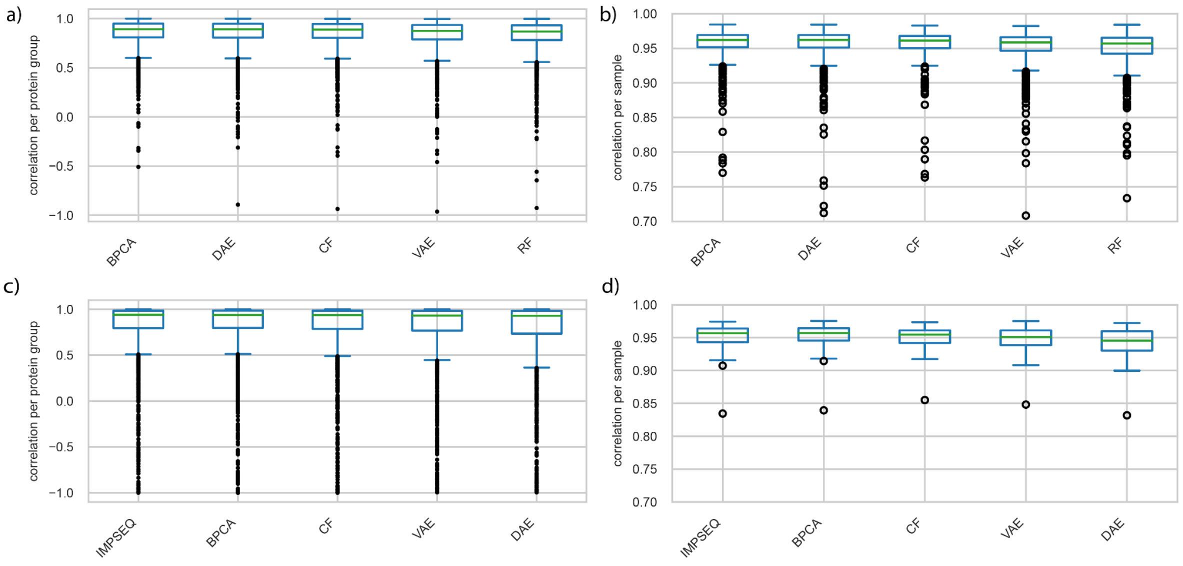


**Fig. S6: Boxplots of correlations between protein groups.** a) Correlation between 4,494 protein groups with at least 3 intensities in test data split with 25 percent MNAR for large development dataset and c) 1,566 protein groups for small development dataset. b) Correlations between protein groups between 564 samples and d) 50 samples. Mean correlation lies around 0.96 for all methods. See **Supp. Data 6** for summary statistics.

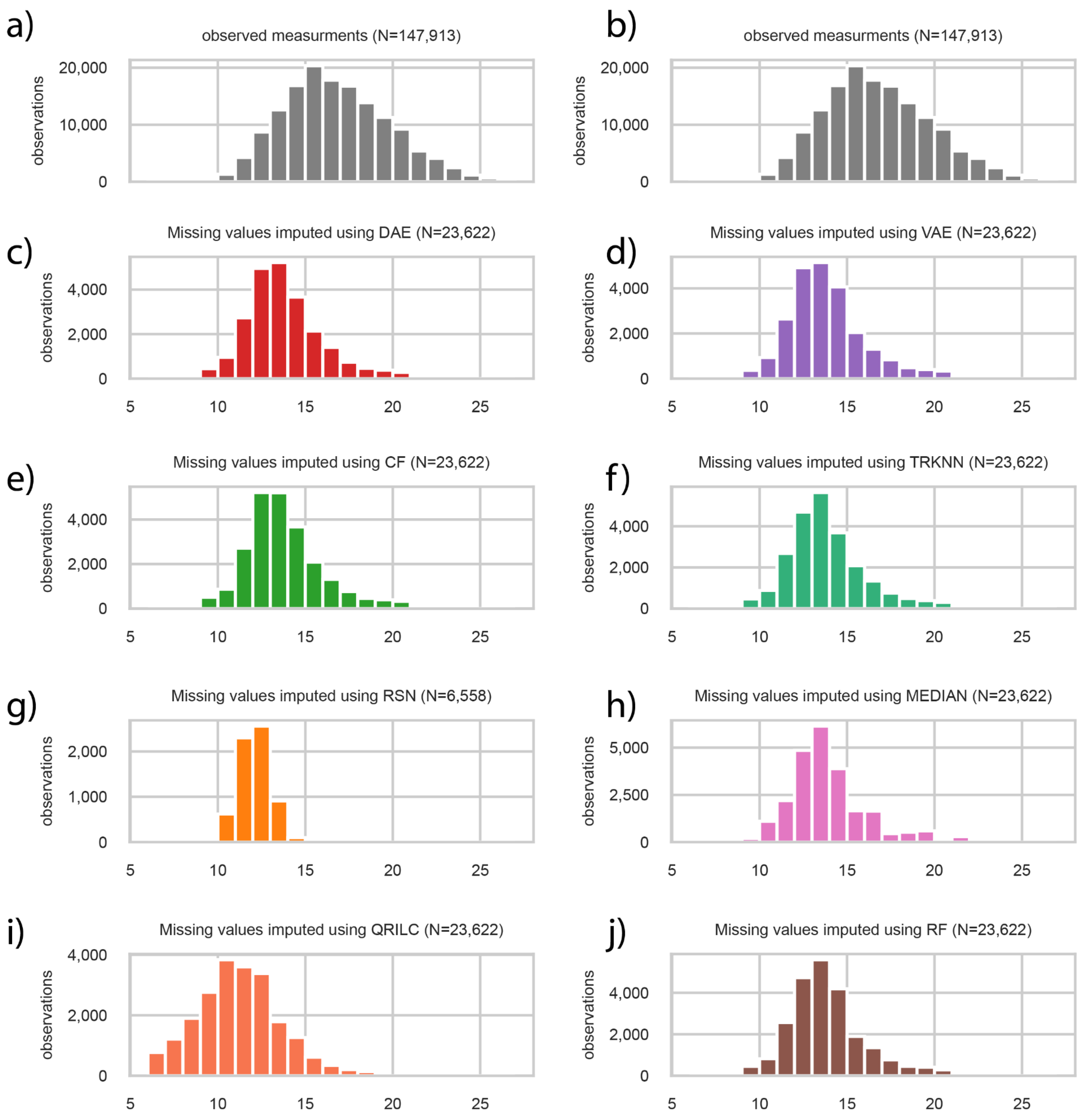


**Fig. S7: ALD plasma protein groups imputations. a-b)** Histogram of all quantified protein group measurements which were available in gray, displayed twice in the first row. **c-j)** Histograms of all model based imputations. **g)** RSN was only used to impute a subset of the features as in the original study. All histogram bars overlap exactly.

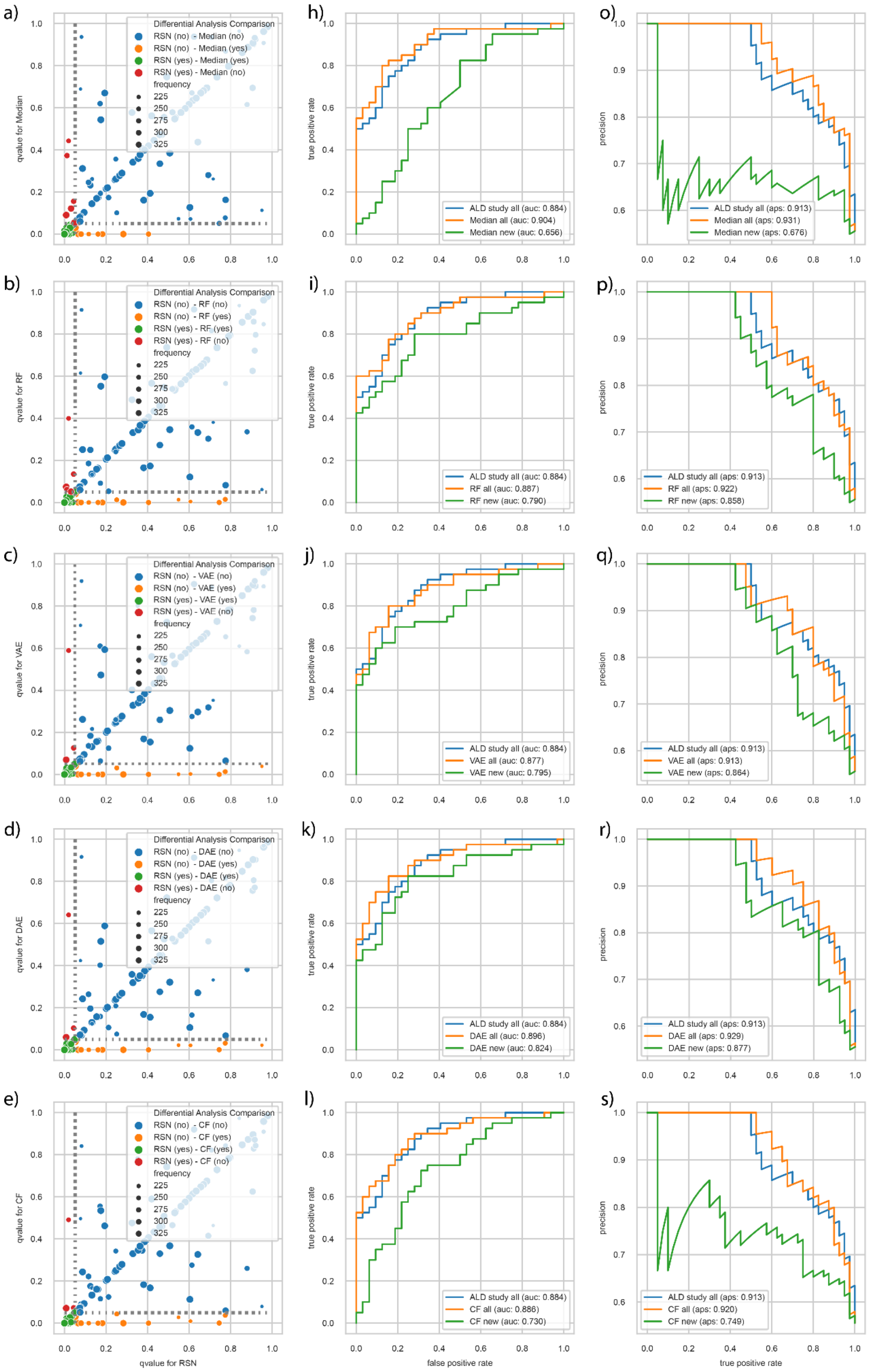

a)
Differential Analysis Comparison
RSN (no) - Median (no)
RSN (no) - Median (yes)
RSN (yes) - Median (yes)
RSN (yes) - Median (no)
frequency
225
250
275
300
325
qvalue for Median
h)
true positive rate
ALD study all (auc: 0.884)
Median all (auc: 0.904)
Median new (auc: 0.656)
o)
precision
ALD study all (aps: 0.913)
Median all (aps: 0.931)
Median new (aps: 0.676)
b)
Differential Analysis Comparison
RSN (no) - RF (no)
RSN (no) - RF (yes)
RSN (yes) - RF (yes)
RSN (yes) - RF (no)
frequency
225
250
275
300
325
qvalue for RF
i)
ALD study all (auc: 0.884)
RF all (auc: 0.887)
RF new (auc: 0.790)
p)
ALD study all (aps: 0.913)
RF all (aps: 0.922)
RF new (aps: 0.858)
c)
Differential Analysis Comparison
RSN (no) - VAE (no)
RSN (no) - VAE (yes)
RSN (yes) - VAE (yes)
RSN (yes) - VAE (no)
frequency
225
250
275
300
325
qvalue for VAE
j)
ALD study all (auc: 0.884)
VAE all (auc: 0.877)
VAE new (auc: 0.795)
q)
ALD study all (aps: 0.913)
VAE all (aps: 0.913)
VAE new (aps: 0.864)
d)
Differential Analysis Comparison
RSN (no) - DAE (no)
RSN (no) - DAE (yes)
RSN (yes) - DAE (yes)
RSN (yes) - DAE (no)
frequency
225
250
275
300
325
qvalue for DAE
k)
ALD study all (auc: 0.884)
DAE all (auc: 0.896)
DAE new (auc: 0.824)
r)
ALD study all (aps: 0.913)
DAE all (aps: 0.929)
DAE new (aps: 0.877)
e)
Differential Analysis Comparison
RSN (no) - CF (no)
RSN (no) - CF (yes)
RSN (yes) - CF (yes)
RSN (yes) - CF (no)
frequency
225
250
275
300
325
qvalue for CF
qvalue for RSN
l)
ALD study all (auc: 0.884)
CF all (auc: 0.886)
CF new (auc: 0.730)
false positive rate
s)
ALD study all (aps: 0.913)
CF all (aps: 0.920)
CF new (aps: 0.749)
true positive rate

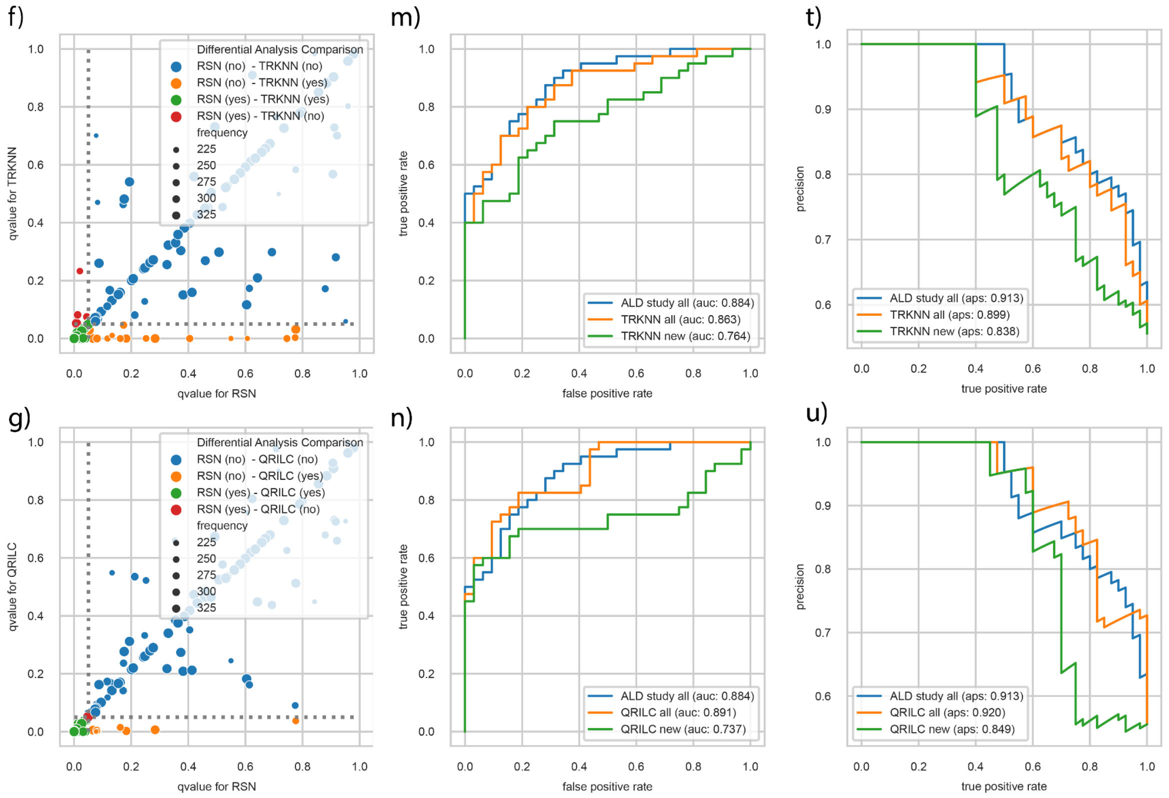


**Fig. S8: Differential analysis comparisons and logistic regression performance using models. a-g)** Comparing qvalues based on RSN imputation (x-axis) to the ones based on median, RF, DAE, VAE, CF, TRKNN and QRILC imputations (y-axis) for the 313 protein groups included in Niu. et al 2021. **h-n)** AUROC of logistic regression on test data allowing selection of original ALD study included 313 protein groups and RSN imputation (blue) to all 377 (orange) or only the 64 new protein groups included here (green). **o-u)** As for h-n showing PRAUC curves for the logistic regression on test data.

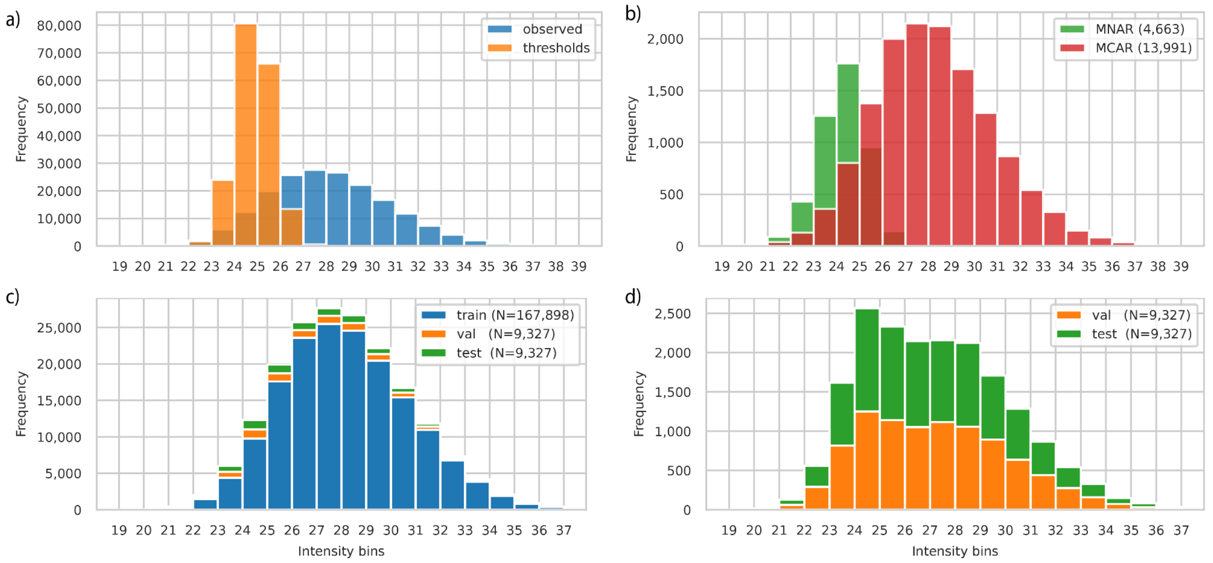


**Fig. S9: Stacked histogram of train, validation and test split for protein groups of smaller development dataset with a share of 25 percent MNAR in the simulated missing values. a)** Histogram of thresholds (orange) for each observed (blue) value for MNAR selection procedure. **b)** Sampled MNAR and MCAR simulated missing values. **c)** Stack histogram of all three splits. **d)** Validation and test data split stacked. Bins were created per integer of logarithm two transformed intensities. Number of intensities per split in parentheses.

### Supplementary Tables

| Method | Package | Source | Description |
|---|---|---|---|
| CF | pimms | PyPI | Collaborative Filtering |
| DAE | pimms | PyPI | Denoising Autoencoder |
| VAE | pimms | PyPI | Variational Autoencoder |
| KNN | Scikit-learn | PyPI | K nearest neighbor imputation |
| RSN | pimms | PyPi | Downshifted normal distribution (per sample), as PI |
| Median | pimms | PyPI | Imputation by feature median |
| ZERO | - | - | replace NA with 0 |
| MINIMUM | - | - | replace NA with global minimum |
| COLMEDIAN | e1071 | CRAN | replace NA with column median |
| ROWMEDIAN | e1071 | CRAN | replace NA with row median |
| KNN_IMPUTE | impute | BIOCONDUCTOR | k nearest neighbor imputation |
| SEQKNN | SeqKnn | tar file | Sequential k- nearest neighbor imputation starts with feature with least missing values and re-use imputed values for not yet imputed features |
| BPCA | pcaMethods | BIOCONDUCTOR | Bayesian PCA missing value imputation |
| SVDMETHOD | pcaMethods | BIOCONDUCTOR | replace NA initially with zero, use k most significant eigenvalues using Singular Value Decomposition for imputation until convergence |
| LLS | pcaMethods | BIOCONDUCTOR | Local least squares imputation of a feature based on k most correlated features |
| MLE | norm | CRAN | Maximum likelihood estimation |
| QRILC | imputeLCMD | CRAN | quantile regression imputation of left-censored data, i.e. by random draws from a truncated distribution which parameters were estimated by quantile regression |
| MINDET | imputeLCMD | CRAN | replace NA with q-quantile minimum in a sample |
| MINPROB | imputeLCMD | CRAN | replace NA by random draws from q-quantile minimum centered distribution |
| IRM | VIM | CRAN | iterativ robust model-based imputation (one feature at at time) |
| IMPSEQ | rrcovNA | CRAN | Sequential imputation of missing values by minimizing the determinant of the covariance matrix with imputed values |
| IMPSEQROB | rrcovNA | CRAN | Sequential imputation of missing values using robust estimators |
| MICE-NORM | mice | CRAN | Multivariate Imputation by Chained Equations (MICE) using Bayesian linear regression |
| MICE-CART | mice | CRAN | Multivariate Imputation by Chained Equations (MICE) using regression trees |
| TRKNN | - | script | truncation k-nearest neighbor imputation |
| RF | missForest | CRAN | Random Forest imputation (one feature at a time) |
| PI | - | - | Downshifted normal distribution (per sample) |
| GSIMP | - | script | QRILC initialization and iterative Gibbs sampling with generalized linear models (glmnet) |
| MSIMPUTE | msImpute | BIOCONDUCTOR | Missing at random algorithm using low rank approximation |
| MSIMPUTE MNAR | msImpute | BIOCONDUCTOR | Missing not at random algorithm using low rank approximation |

**Table S1**: **Methods overview in current comparison**. Sources for packages are mainly PyPI for PIMMS, CRAN and Bioconductor for R packages. Some R packages require custom installations. Find the up-to-date overview on GitHub.

| Gene | Freq. observed | CF | DAE | Median | None | QRILC | RF | TRKNN | VAE |
|---|---|---|---|---|---|---|---|---|---|
| S100A9 | 168 | 10 | 10 | 0 | 0 | 0 | 10 | 10 | 10 |
| PEBP4 | 124 | 10 | 10 | 0 | 0 | 0 | 10 | 10 | 10 |
| GLIPR2 | 167 | 10 | 10 | 0 | 0 | 3 | 10 | 10 | 10 |
| IGKV1-37 | 156 | 10 | 10 | 0 | 0 | 9 | 10 | 10 | 10 |
| HSPG2 | 137 | 10 | 1 | 10 | 0 | 10 | 7 | 10 | 5 |
| PRDX2 | 110 | 10 | 10 | 0 | 0 | 4 | 9 | 10 | 10 |
| KRT25 | 148 | 3 | 9 | 0 | 0 | 0 | 0 | 10 | 7 |
| RPS6KA3 | 179 | 0 | 0 | 0 | 0 | 4 | 0 | 0 | 0 |
| TLN1 | 106 | 0 | 3 | 0 | 0 | 0 | 1 | 0 | 9 |
| CCT3 | 108 | 0 | 2 | 0 | 0 | 0 | 0 | 0 | 2 |
| ALDOA | 102 | 10 | 10 | 0 | 0 | 0 | 8 | 10 | 10 |
| ANK1 | 112 | 10 | 10 | 0 | 0 | 8 | 10 | 10 | 10 |
| RAB2B | 187 | 10 | 10 | 0 | 0 | 0 | 10 | 10 | 10 |
| PZP | 200 | 10 | 4 | 0 | 0 | 0 | 7 | 10 | 10 |
| HSPA1B;HSPA1A | 107 | 10 | 10 | 0 | 0 | 0 | 10 | 10 | 10 |
| AMY1C | 92 | 10 | 10 | 10 | 0 | 7 | 10 | 10 | 9 |
| ALCAM | 124 | 10 | 10 | 0 | 0 | 10 | 10 | 10 | 10 |
| CCT6A | 139 | 0 | 2 | 0 | 0 | 0 | 0 | 0 | 0 |
| GSTO1 | 111 | 0 | 1 | 0 | 0 | 3 | 0 | 10 | 0 |
| C16orf46 | 195 | 0 | 1 | 0 | 0 | 0 | 0 | 0 | 0 |
| CA3 | 152 | 0 | 8 | 0 | 0 | 0 | 1 | 10 | 7 |
| P4HB | 111 | 0 | 3 | 0 | 0 | 0 | 3 | 10 | 2 |
| REG1A;REG1B | 107 | 7 | 0 | 0 | 0 | 1 | 6 | 10 | 3 |
| FLNA | 99 | 10 | 9 | 0 | 0 | 0 | 9 | 10 | 9 |
| ZNF511-PRAP1;PRAP1 | 148 | 0 | 5 | 0 | 0 | 0 | 4 | 10 | 6 |
| SOD3 | 117 | 0 | 0 | 0 | 0 | 0 | 0 | 10 | 0 |
| TMSB4X | 125 | 0 | 0 | 0 | 0 | 0 | 0 | 10 | 0 |
| DPP4 | 110 | 0 | 0 | 0 | 0 | 0 | 0 | 10 | 1 |
| ICAM2 | 132 | 10 | 10 | 0 | 0 | 10 | 10 | 10 | 9 |
| KRT10 | 117 | 0 | 1 | 0 | 0 | 0 | 0 | 10 | 1 |
| DCD | 94 | 0 | 0 | 0 | 0 | 0 | 0 | 10 | 0 |
| RARRES2 | 110 | 0 | 2 | 0 | 0 | 0 | 1 | 10 | 6 |
| ITSN1 | 124 | 10 | 8 | 0 | 0 | 2 | 1 | 0 | 4 |
| SAA2 | 130 | 10 | 8 | 0 | 0 | 0 | 10 | 0 | 5 |
| APMAP | 196 | 0 | 0 | 0 | 0 | 2 | 0 | 10 | 0 |
| NPEPPS | 180 | 0 | 0 | 0 | 0 | 1 | 0 | 0 | 0 |
| PPIA | 185 | 0 | 0 | 0 | 0 | 2 | 0 | 0 | 0 |
| FAH | 115 | 0 | 1 | 0 | 0 | 0 | 0 | 0 | 1 |

**Table S2: Counts of how often 54 differentially abundant protein groups which were newly included were found by any included model to be significant running the analysis 10 times.** No imputation (None), Median and TRKNN imputation did not change between runs (**Supp. Data. 12**).
